# *Sphingobium* sp. SYK-6 syringate *O*-demethylase gene is regulated by DesX, unlike other vanillate and syringate catabolic genes regulated by DesR

**DOI:** 10.1101/2020.07.27.224295

**Authors:** Takuma Araki, Kenta Tanatani, Naofumi Kamimura, Yuichiro Otsuka, Muneyoshi Yamaguchi, Masaya Nakamura, Eiji Masai

**Author notes:** Address correspondence to Eiji Masai.

## Abstract

Syringate and vanillate are the major metabolites of lignin biodegradation. In *Sphingobium* sp. strain SYK-6, syringate is O demethylated to gallate by consecutive reactions catalyzed by DesA and LigM, and vanillate is O demethylated to protocatechuate by a reaction catalyzed by LigM. The gallate ring is cleaved by DesB, and protocatechuate is catabolized via the protocatechuate 4,5-cleavage pathway. The transcriptions of *desA, ligM*, and *desB* are induced by syringate and vanillate, while that of *ligM* and *desB* are negatively regulated by the MarR-type transcriptional regulator DesR, which is not involved in *desA* regulation. Here we clarified the regulatory system for *desA* transcription by analyzing the IclR-type transcriptional regulator *desX*, located downstream of *desA*. Quantitative reverse transcription (RT)-PCR analyses of a *desX* mutant indicates that the transcription of *desA* was negatively regulated by DesX. In contrast, DesX was not involved in the regulation of *ligM* and *desB*. The ferulate catabolic genes (*ferBA*) under the control of a MarR-type transcriptional regulator FerC are located upstream of *desA*. RT-PCR analyses suggest that the *ferB*-*ferA*-SLG_25010-*desA* gene cluster consists of the *ferBA* operon and the SLG_25010-*desA* operon. Promoter assays reveal that a syringate- and vanillate-inducible promoter is located upstream of SLG_25010. Purified DesX bound to this promoter region, which overlaps with an 18-bp-inverted repeat sequence that appears to be essential for the DNA binding of DesX. Syringate and vanillate inhibited the DNA binding of DesX, indicating that these compounds are effector molecules of DesX.

**IMPORTANCE:** Syringate is a major degradation product in the microbial and chemical degradation of syringyl lignin. Along with other low-molecular-weight aromatic compounds, syringate is produced by chemical lignin depolymerization. Converting this mixture into value-added chemicals using bacterial metabolism (i.e., biological funneling) is a promising option for lignin valorization. To construct an efficient microbial lignin conversion system, it is necessary to identify and characterize the genes involved in the uptake and catabolism of lignin-derived aromatic compounds and elucidate their transcriptional regulation. In this study, we found that the transcription of *desA*, encoding syringate *O*-demethylase in SYK-6, is regulated by an IclR-type of transcriptional regulator, DesX. The findings of this study, combined with our previous results on *desR* (a MarR transcriptional regulator that controls the transcription of *ligM* and *desB*), provide an overall picture of the transcriptional regulatory systems for syringate and vanillate catabolism in SYK-6.

## INTRODUCTION

Lignin, a major component of plant cell walls, is the most abundant aromatic polymer in nature, thus its decomposition is essential for the Earth’s carbon cycle (1). Lignin is generated by the oxidative coupling of three types of *p*-hydroxyphenylpropanoids, called monolignols: coniferyl alcohol, sinapyl alcohol, and *p*-coumaryl alcohol (2). Lignin structure differs depending on the type of plant; gymnosperm (softwood) lignins mainly contain guaiacyl (G)-units derived from coniferyl alcohol; angiosperm (hardwood) lignins mainly contain syringyl (S)-units derived from sinapyl alcohol and G-units; and monocot (grass) lignins contain G-units, S-units, and H-units derived from *p*-coumaryl alcohol (3, 4). Because lignin is abundant and renewable, it holds great potential as a bioresource. One option to realize this potential is through biological funneling, a process that uses bacterial catabolic functions to transform heterogeneous mixtures of low-molecular-weight aromatic compounds produced by chemical lignin depolymerization into valuable platform chemicals (5-7).

*Sphingobium* sp. strain SYK-6 is an aerobic alphaproteobacterium with the best-characterized catabolic systems for lignin-derived aromatic compounds (7, 8). As more details on the SYK-6 catabolic system and its regulation emerge, our understanding of bacterial lignin catabolism is enhanced, and such information can also be used to transform lignin through biological funneling (7-11). SYK-6 is capable of utilizing various types of lignin-derived biaryls (e.g., β-aryl ether, biphenyl, phenylcoumaran, and diarylpropane) and monoaryls (e.g., ferulate, vanillin, and syringaldehyde) as its sole carbon and energy source (7, 8). These aromatic compounds, derived from G- and S-units, are degraded through vanillate (VA) and syringate (SA), respectively. Both are common intermediate metabolites in the biodegradation of lignin. VA is then converted to protocatechuate (PCA) by tetrahydrofolate (H_4_folate)-dependent VA/3-*O*-methyl gallate (3MGA) *O*-demethylase (LigM). The resulting PCA is further metabolized via the PCA 4,5-cleavage pathway that includes a step catalyzed by PCA 4,5-dioxygenase (LigAB) (Fig. 1) (12). On the other hand, SA is converted to 3MGA by H_4_folate-dependent SA *O*-demethylase (DesA), whose sequence is 49% similar to LigM in SYK-6 (Fig. 1). The resulting 3MGA is metabolized by branching into three pathways; among them, the gallate (GA) cleavage pathway that is involved crucially in the process includes O demethylation by LigM and ring-cleavage by GA dioxygenase (DesB) (Fig. 1) (13).

**Fig. 1.**
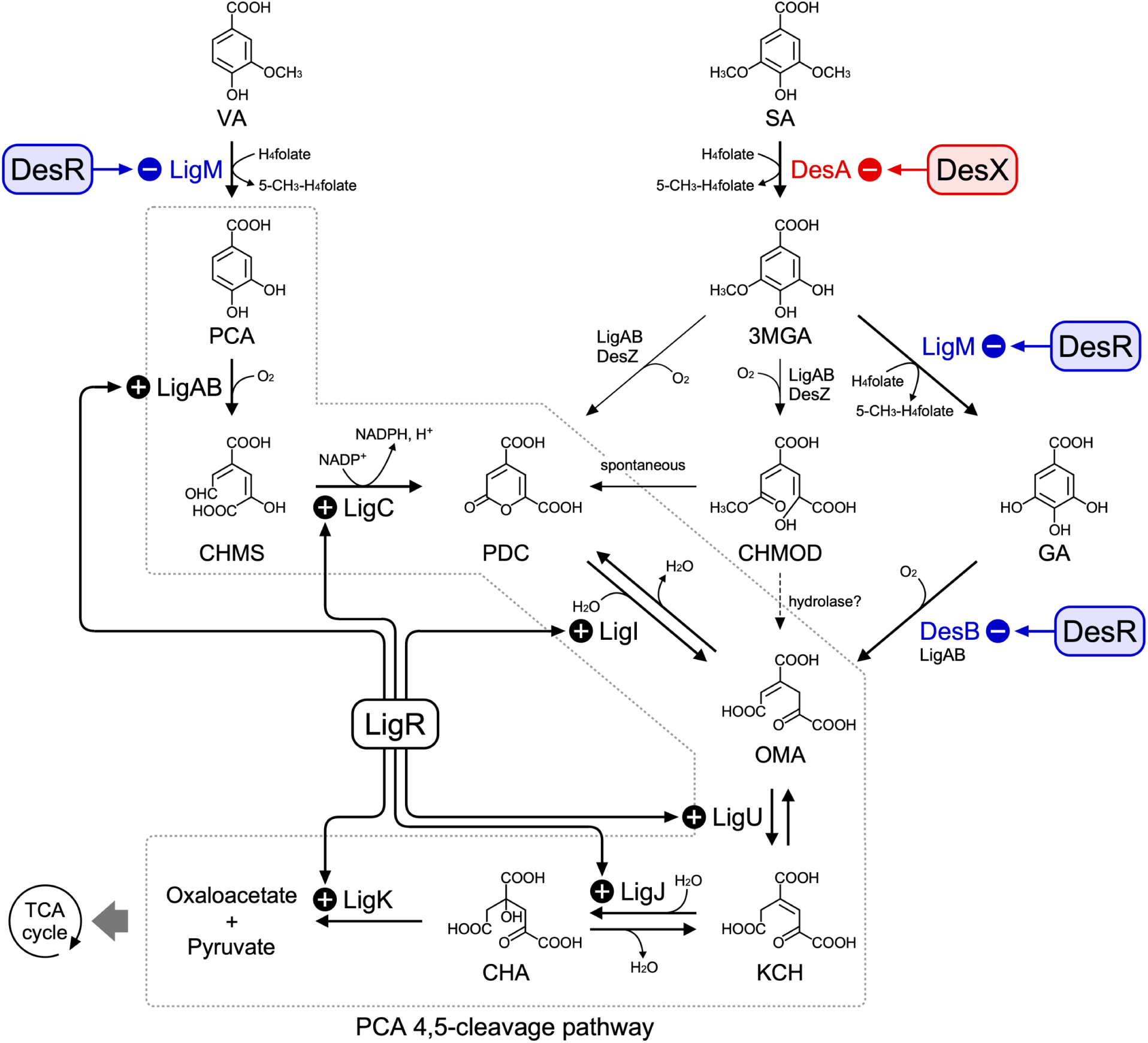
Transcriptional regulation of catabolism of vanillate and syringate in *Sphingobium* sp. SYK-6. The transcriptional regulation of *ligM* and *desB* by DesR (27) and the regulation of the PCA 4,5-cleavage genes by LigR (26) are highlighted in blue background and black bold, respectively. The transcriptional regulation of *desA* by DesX investigated in this study is highlighted by the red background. Enzymes: LigM, vanillate/3MGA *O*-demethylase; LigA and LigB, small and large subunits, respectively, of PCA 4,5-dioxygenase; LigC, CHMS dehydrogenase; LigI, PDC hydrolase; LigU, OMA delta-isomerase; LigJ, KCH hydratase; LigK, CHA aldolase; DesA, syringate *O*-demethylase; DesZ, 3MGA 3,4-dioxygenase; DesB, gallate dioxygenase. Transcriptional regulators: DesX, IclR-type regulator; DesR, MarR-type regulator; LigR, LysR-type regulator. Abbreviations: VA, vanillate; PCA, protocatechuate; CHMS, 4-carboxy-2-hydroxymuconate-6-semialdehyde; PDC, 2-pyrone-4,6-dicarboxylate; OMA, 4-oxalomesaconate; KCH, 2-keto-4-carboxy-3-hexenedioate; CHA, 4-carboxy-4-hydroxy-2-oxoadipate; SA, syringate; 3MGA, 3-*O*-methylgallate; GA, gallate; CHMOD, 4-carboxy-2-hydroxy-6-methoxy-6-oxohexa-2,4-dienoate.

The bacterial degradation of VA through the pathway composed of an oxygenase-type VA *O*-demethylase (VanAB) and the PCA 3,4-cleavage pathway enzymes has been mainly investigated in *Pseudomonas* strains (14, 15), *Acinetobacter baylyi* ADP1 (16), *Caulobacter crescentus* (17, 18), *Corynebacterium glutamicum* ATCC 13032 (19), and *Rhodococcus jostii* RHA1 (20). In *A. baylyi* ADP1, *C. crescentus*, and *C. glutamicum* ATCC 13032, the transcriptional regulation of *vanAB* is negatively regulated by a GntR-type, GntR-type, and PadR-like transcriptional regulator, respectively, all of which are called VanR (17, 21, 22). VA was determined to be an effector molecule for the latter two systems (17, 22). The degradation of SA by bacteria other than SYK-6 has recently been reported in *Novosphingobium aromaticivorans* DSM 12444 (23), *Microbacterium* sp. strain RG1 (24), and *Pseudomonas* sp. strain NGC7 (25). While the SA catabolic pathway genes have been identified in DSM 12444 and predicted in RG1, the regulatory system of SA catabolism has not been studied in any bacterium.

In terms of the transcriptional regulation involved in VA and SA catabolism in SYK-6, we have reported that the PCA 4,5-cleavage pathway genes are positively regulated by LigR, a LysR-type transcriptional regulator that recognizes PCA and GA as effectors (Fig. 1) (26). Moreover, we have shown that a MarR-type transcriptional regulator, DesR, negatively regulates the transcription of *ligM* and *desB*, and VA and SA are effectors that release the repression by DesR (Fig. 1) (27). Although the transcription of *desA* is also induced by VA and SA, DesR does not participate in the transcriptional regulation of *desA*; thus, its regulatory system remains unknown.

In this study, we clarified the regulatory system of SA catabolism in SYK-6 by identifying and characterizing an IclR-type transcriptional regulator, DesX, which regulates *desA* transcription.

## RESULTS

### Search for genes involved in transcriptional regulation of *desA*

To determine the presence of a transcriptional regulator of *desA* in *Sphingobium* sp. SYK-6, we performed an electrophoretic mobility shift assay (EMSA) using SYK-6 cell extracts and desAp1−desAp5 probes containing sequences upstream of, and within *desA*, as shown in Fig. 2A. SYK-6 was grown in Wx minimal medium (28) containing 10 mM sucrose, 10 mM glutamate, 0.13 mM methionine, and 10 mM proline (Wx-SEMP; the noninducing conditions) and in Wx-SEMP containing 5 mM VA or SA (the VA/SA-inducing conditions). The cell extracts from these cultures were incubated with each of the desAp1−desAp5 probes. A band shift was observed only with desAp2, regardless of the cell extracts used (Fig. 2B). These results indicate that an unknown SYK-6 protein binds to the DNA region spanning from 103-bp upstream to 229-bp downstream of the initiation codon of SLG_25010 (which is just upstream of *desA*).

**Fig. 2.**
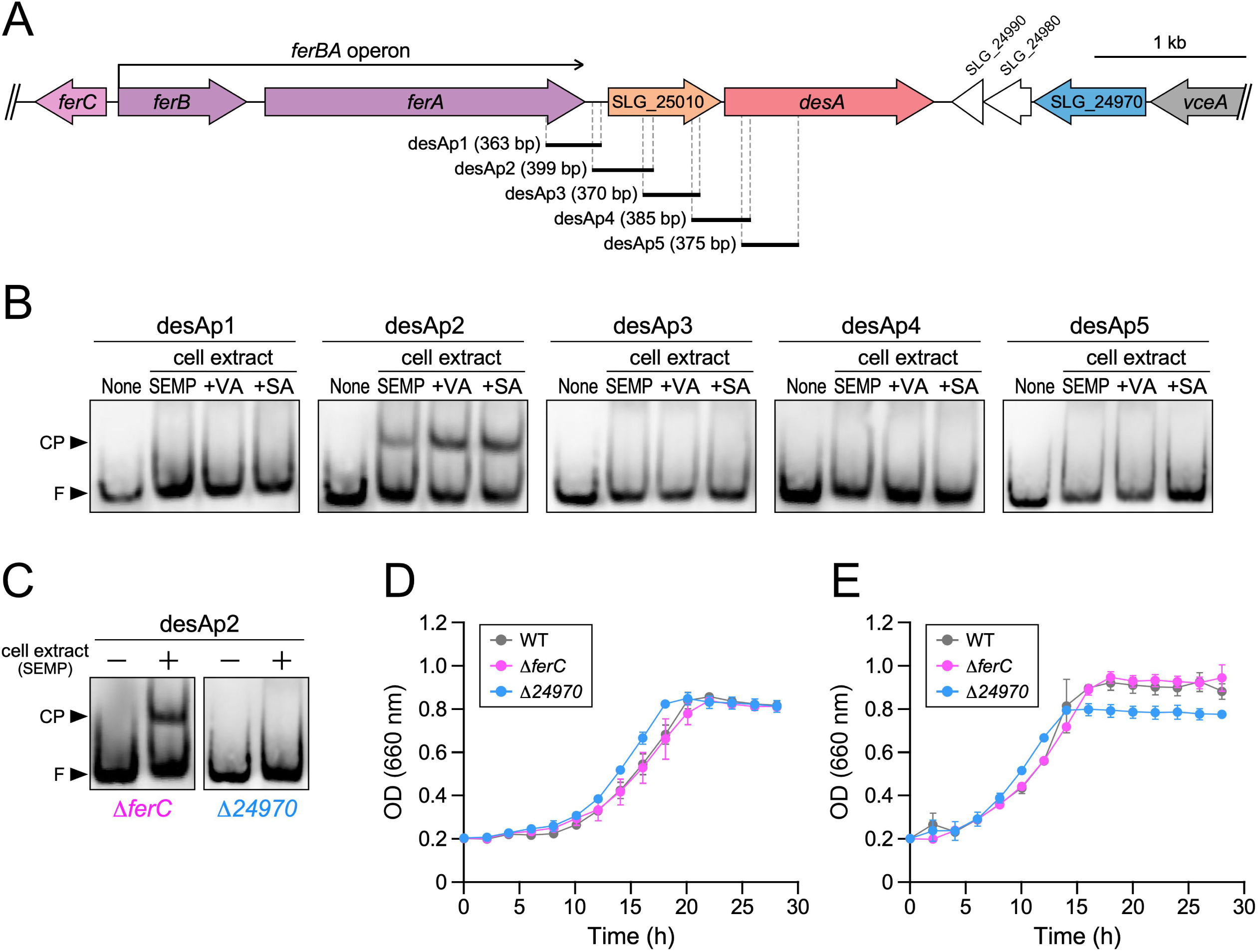
Identification of an SYK-6 protein that binds to the upstream region of *desA*. (A) Gene organization around *desA*. Genes: *ferC*, MarR-type transcriptional regulator; *ferB*, feruloyl-CoA hydratase/lyase; *ferA*, feruloyl-CoA synthetase; SLG_25010, putative hydrolase; SLG_24970 (*desX*), IclR-type transcriptional regulator; *vceA*, vanilloyl acetic acid/3-(4-hydroxy-3,5-dimethoxyphenyl)-3-oxopropanoic acid-converting enzyme (49). Black bars under the map show the DNA fragments used for EMSA (desAp1−desAp5 probes). (B) EMSAs of SYK-6 cell extracts using the desAp1−desAp5 probes. Digoxigenin-labeled probes (500 pM) were incubated in the presence and absence of the extracts (0.4 μg protein/μl) of wild type cells grown in Wx-SEMP, Wx-SEMP + VA, and Wx-SEMP + SA. CP, DNA-protein complex; F, free probe. (C) EMSAs of Δ*ferC* and Δ*24970* cell extracts using the desAp2 probe. The desAp2 probe (500 pM) was incubated in the presence (+) and absence (−) of the extracts (0.4 μg protein/μl) of Δ*ferC* and Δ*24970* cells grown in Wx-SEMP. (D and E) Growth of Δ*ferC* and Δ*24970* on SA or VA, respectively. Cells of SYK-6 (gray), Δ*ferC* (magenta), and Δ*24970* (cyan) were incubated in Wx-5 mM SA (D) or Wx-5 mM VA (E), and OD_660_ was periodically monitored. Each value is the average ± the standard deviation (error bars) of three independent experiments.

Upstream of *desA* (SLG_25000) are *ferC* (SLG_25040) and the *ferBA* (SLG_25030-25020) operon involved in ferulate (FA) catabolism. The *ferC* gene encodes a MarR-type transcriptional regulator that negatively regulates *ferBA* (Fig. 2A). Downstream of *desA*, there is SLG_24970, which is similar to the IclR-type transcriptional regulator (ITTR) (Fig 2A; Table S1). To clarify whether the gene product of *ferC* or SLG_24970 is involved in the binding of the desAp2 probe, we conducted EMSAs using cell extracts of a *ferC* mutant (Δ*ferC*) and an SLG_24970 mutant (Δ*24970*). Δ*ferC* was obtained in our previous study (28), while Δ*24970* was constructed through homologous recombination in this study (Fig. S1). EMSAs using a cell extract of Δ*ferC* grown in Wx-SEMP show a band shift similar to that of the wild type; however, no such band shift occurs in EMSA using a cell extract of Δ*24970* grown in the same medium (Fig. 2C). These results strongly suggest that the band shift was due to the binding of the SLG_24970 gene product to the desAp2 region.

To determine whether the disruption of *ferC* and SLG_24970 affects SA and VA catabolism in SYK-6, we measured the growth of Δ*ferC* and Δ*24970* on 5 mM SA and VA. Δ*24970* grew on SA and VA somewhat faster than the wild type, while Δ*ferC* grew on SA and VA as well as the wild type (Fig. 2D and E). These results suggest that the SLG_24970 gene product negatively regulates the transcription of *desA* by binding to the desAp2 region. Introduction of a plasmid carrying SLG_24970 (pJB24970) into Δ*24970* and SYK-6 caused a substantial delay in their growth on SA compared to Δ*24970* and SYK-6 harboring the vector (pJB866) (Fig. S2). These results indicate that the disruption of SLG_24970 caused the changed phenotype of Δ*24970*. Thus, we designated SLG_24970 as *desX*.

### Transcriptional analysis of the SA and VA catabolic genes in a *desX* mutant

To determine whether *desX* is involved in the transcriptional regulation of the SA and VA catabolic genes *desA, ligM*, and *desB* in SYK-6, quantitative reverse transcription-PCR (qRT-PCR) analyses of these genes were performed using total RNAs isolated from the wild type and Δ*desX* cells grown under the noninducing and the VA/SA-inducing conditions. The transcription levels of *desA, ligM*, and *desB* in the wild type under VA/SA-inducing conditions increased 28- to 50-, 5.7- to 14-, and 4.4- to 7.6-fold, respectively, compared to noninducing conditions (Fig. 3A to C). In Δ*desX* cells under noninducing conditions, the transcription levels of *ligM* and *desB* were similar to those of the wild type under the same conditions. Additionally, the transcription of *ligM* and *desB* was activated in Δ*desX* cells as well as in the wild type under VA/SA-inducing conditions (Fig. 3B and C). By contrast, the transcription levels of *desA* in Δ*desX* cells under noninducing conditions increased 48-fold compared to that of the wild type under the same conditions, and were comparable to that of the wild type under inducing conditions (Fig. 3A). Therefore, it is evident that the transcription of *desA* is negatively regulated by DesX. The *desA* transcription levels were close to that of Δ*desX* cells under both noninducing and VA/SA-inducing conditions. These results suggest that the regulation of *desA* transcription is governed by DesX, and DesX is not involved in the transcriptional regulation of *ligM* and *desB*.

**Fig. 3.**
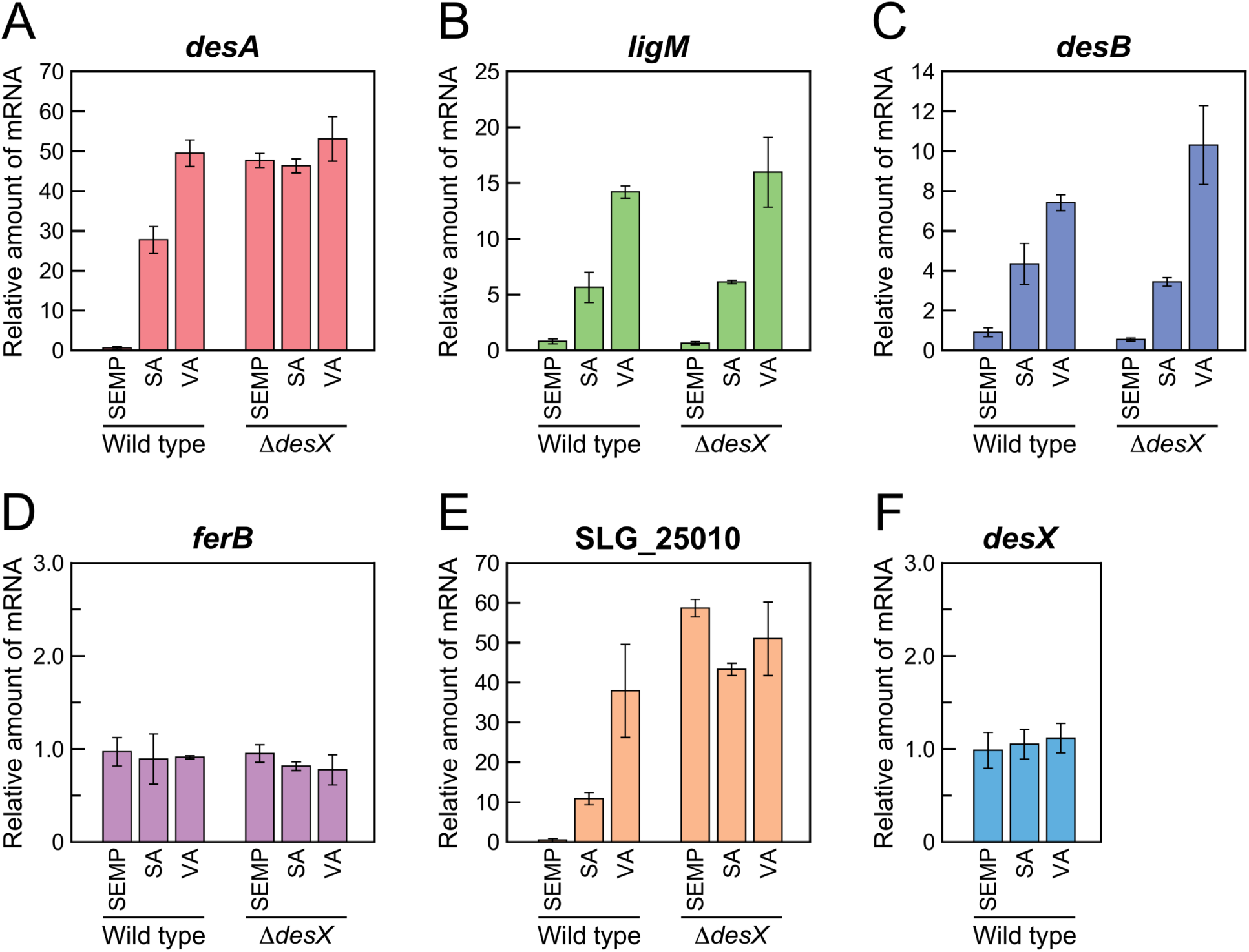
qRT-PCR analysis of the expression of *desA, ligM, desB, ferB*, SLG_25010, and *desX* (SLG_24970). Total RNAs were isolated from the cells of SYK-6 and a *desX* mutant (Δ*desX*) grown in Wx-SEMP, Wx-5 mM SA, and Wx-5 mM VA. The relative mRNA amounts of *desA* (A), *ligM* (B), *desB* (C), *ferB* (D), SLG_25010 (E), and *desX* (F; measured only in the wild type) indicate the fold increases relative to the amount of mRNA in SYK-6 cells grown in Wx-SEMP (level of 1.0). Values for each amount of mRNA were normalized to the level of 16S rRNA. Each value is the average ± the standard deviation (error bars) of three independent experiments.

Upstream of *desA* is SLG_25010, which encodes a putative hydrolase and the *ferBA* operon both of which are transcribed in the same direction as *desA* (Fig. 2A). qRT-PCR analyses show that the transcription levels of *ferB* in the wild type and Δ*desX* cells were almost the same under both noninducing and SA/VA-inducing conditions (Fig. 3D), indicating that DesX does not participate in the regulation of the *ferBA* operon. By contrast, the transcription of SLG_25010 in the wild type increased 11- to 39-fold under SA/VA-inducing conditions compared to noninducing conditions (Fig. 3E). In Δ*desX* cells under noninducing conditions, the transcription level of SLG_25010 increased 59-fold compared to that of the wild type grown under the same conditions. This level is equivalent to that in Δ*desX* cells under SA/VA-inducing conditions (44- to 51-fold) (Fig. 3E). Therefore, in the transcriptional regulation of SLG_25010, DesX acts as a repressor, while SA and VA appear to be inducers of SLG_25010, along with *desA*. In addition, the transcription level of *desX* in the wild type remained nearly constant regardless of the culture conditions (Fig. 3F), indicating that *desX* is constitutively expressed.

### Operon structure of the *ferB*-*ferA*-SLG_25010-*desA* gene cluster

In our previous study, the *ferB*-*ferA*-SLG_25010-*desA* gene cluster formed an operon during growth in the presence of SA (29). By contrast, although *ferB* and *ferA* formed an operon during growth in the presence of FA, SLG_25010 was not co-transcribed with *ferBA*. In this study, qRT-PCR analysis indicates that the transcriptional inducibility of *ferBA* and SLG_25010-*desA* in SYK-6 cells grown on SA and VA were completely different (Fig. 3). We, therefore, reevaluated the operon structure of the *ferB*-*ferA*-SLG_25010-*desA* gene cluster by RT-PCR analysis using total RNA isolated from SYK-6 grown in Wx-SEMP (the noninducing conditions) and Wx medium containing 5 mM SA, VA, or FA (the SA/VA/FA-inducing conditions) (Fig. S3A).

Under noninducing conditions, only the *ferB*-*ferA* region was amplified. In contrast, under SA/VA/FA-inducing conditions, the *ferA*-SLG_25010 and SLG_25010-*desA* regions were amplified, in addition to the *ferB*-*ferA* region, although the amplification of the *ferA*-*desA* region was not observed (Fig. S3B to E). These results strongly suggest that the *ferB*-*ferA*-SLG_25010-*desA* gene cluster consists of the *ferBA* operon and the SLG_25010-*desA* operon. The amplification of *ferA*-SLG_25010 is likely a consequence of the read-through from the *ferB* promoter. Why was the *ferA*-SLG_25010 region amplified under SA/VA-inducing conditions that does not induce the *ferBA* operon, but not under noninducing conditions? We speculate that when SYK-6 cells were incubated under noninducing conditions, DesX binding to the promoter region of the SLG_25010-*desA* operon probably inhibited the transcription from the *ferB* promoter. By contrast, when SYK-6 cells were incubated under SA/VA-inducing conditions, DesX was released from the promoter region, thus, allowing the transcription from the *ferB* promoter to proceed.

### Determination of the promoter region of the SLG_25010-*desA* operon

Promoter assays using *lacZ* as the reporter gene were performed to determine the promoter region of the SLG_25010-*desA* operon. pSDA1−4 were constructed by cloning DNA fragments containing various regions from *ferA* to *desA* into the region upstream of *lacZ* in a promoter probe vector, pSEVA225 (Fig. S4A). Promoter activities of SYK-6 cells harboring each pSDA1−4 grown under noninducing and SA/VA-inducing conditions were then measured as β-galactosidase activities (Fig. S4B). Prominent promoter activities were observed only in SYK-6 cells harboring pSDA2 under SA/VA-inducing conditions that elicited activities that were 12 times higher than that under noninducing conditions. These results suggest that a SA/VA-inducible promoter is located in the intergenic region between *ferA* and SLG_25010.

The transcription start site of the SLG_25010-*desA* operon was determined by primer extension analysis using a fluorescently labeled oligonucleotide primer and total RNA isolated from SYK-6 cells grown on SA. This analysis yielded a 76-nucleotide extension product that maps the transcription start site of the SLG_25010-*desA* operon to the T residue located 10 nucleotides upstream of the initiation codon of SLG_25010 (Fig. S5). The putative −35 and −10 sequences, bearing similarities to the conserved sequences of the *E. coli* σ^70^-dependent promoter, were found upstream of the transcription start site of the SLG_25010-*desA* operon (Fig. S5). Additionally, we found an 18-bp-inverted repeat sequence, named IR-DA (5’-TCTTCGTATATACGAAGA-3’) at positions −22 to −5 (overlapping the putative −10 sequence) (Fig. S5).

To determine whether the putative −35 and −10 sequences are necessary for the transcription of the SLG_25010-*desA* operon, deletions were introduced in the putative promoter region in pSDA2 (Fig. 4). Like SYK-6 cells harboring pSDA2, SYK-6 cells harboring pSDA2a (containing putative −35 and −10 sequences) exhibited promoter activity that was 9.4-fold higher under SA/VA-inducing conditions compared to activity under noninducing conditions. However, the promoter activity was not observed in SYK-6 cells harboring pSDA2b (containing only putative −10 sequence) or pSDA2c (lacking putative −35 and −10 sequences) (Fig. 4). These results suggest that the putative −35 and −10 sequences function as the essential promoter of the SLG_25010-*desA* operon.

**Fig. 4.**
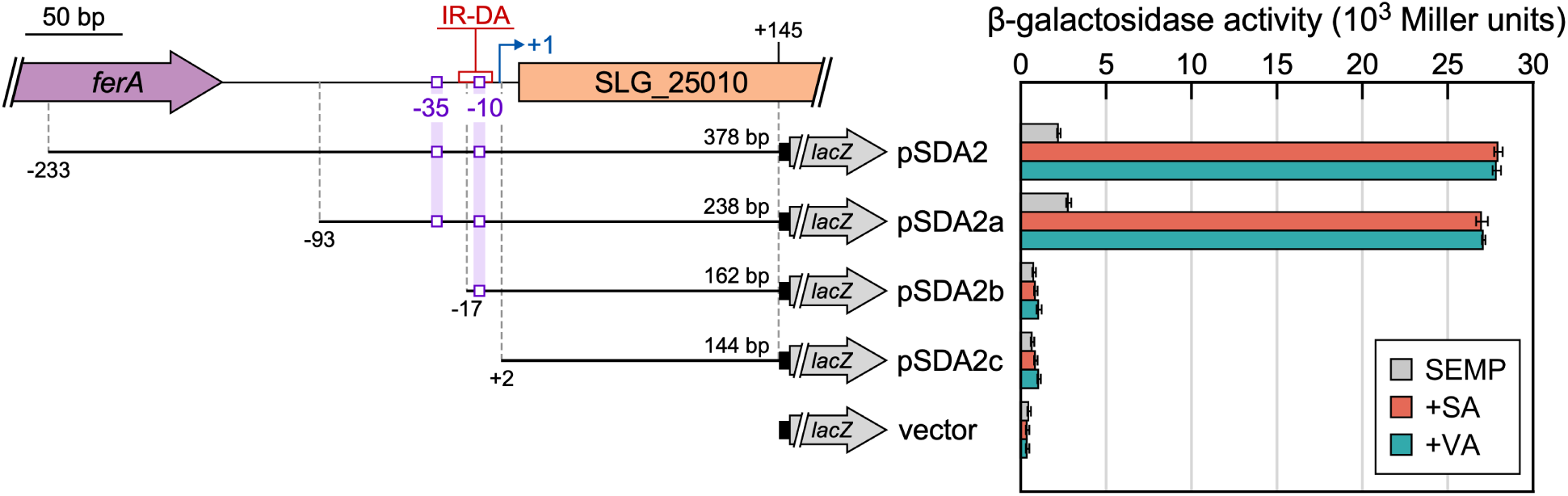
Deletion analysis of the SLG_25010 promoter region. (Left) The DNA fragments used for the promoter analysis of the SLG_25010-*desA* operon. The transcription start site of the SLG_25010-*desA* operon is shown by a bent arrow. Positions of IR-DA and the putative −35 and −10 sequences are indicated. (Right) β-galactosidase activities of SYK-6 cells harboring each reporter plasmid grown in the presence and absence of 5 mM SA or VA. Each value is the average ± the standard deviation (error bars) of three independent experiments.

### Binding of DesX to the promoter region of the SLG_25010-*desA* operon

*desX* fused with a His-tag at the 5’ terminus was expressed under the control of the T7 promoter in *E. coli* BL21(DE3) harboring p16bdX (Table 1). Sodium dodecyl sulfate-polyacrylamide gel electrophoresis (SDS-PAGE) analysis shows the His-tag fused DesX as an approximately 29-kDa protein, which is close to the theoretical molecular weight of 28,968 (Fig. S6). DesX was purified to near homogeneity using Ni affinity chromatography (Fig. S6).

EMSAs were performed to determine whether DesX binds to the promoter region of the SLG_25010-*desA* operon using DNA probes containing various regions around the SLG_25010-*desA* operon (desAp1−desAp5 probes). Probes were incubated with purified DesX (Fig. 5A), resulting in a band shift that shows the formation of a DNA-DesX complex with the desAp2 probe that contains the promoter region of the SLG_25010-*desA* operon. The other DNA probes generated no other band shifts (Fig. 5B). Incubating the desAp2 probe with different concentrations of DesX resulted in one band shift for all concentrations (Fig. S7), implying that there is one binding site for DesX in the desAp2 region. To determine if IR-DA is required for DNA binding of DesX, we performed EMSAs using the desAp6−desAp8 probes shown in Fig. 5A. A band shift was observed only for the desAp7 probe that contains IR-DA (Fig. 5B). We further examined the binding of DesX to DNA probes containing mutated IR-DA. EMSA using the desAp9 fragment containing IR-DA at the 3’ end shows the formation of a DNA-DesX complex (Fig. 5A and C). Using the desAp9mL and desAp9mR probes containing a 9-base mutation in the left and right halves of the desAp9 probe, respectively, resulted in significantly reduced band shifting (Fig. 5A and C). All these results make clear that IR-DA is essential for the DNA binding of DesX. Additionally, we performed EMSAs using DNA probes containing the *ferB, ligM*, and *desB* promoter regions, which include the binding sites of FerC, DesR, and DesR, respectively (Fig. S8). DesX did not bind to any of these DNA probes, regardless of the presence or absence of SA or VA. This supports the results of qRT-PCR analyses, indicating that DesX is not involved in regulating the *ferBA* operon, *ligM*, and *desB*.

**Table 1.**
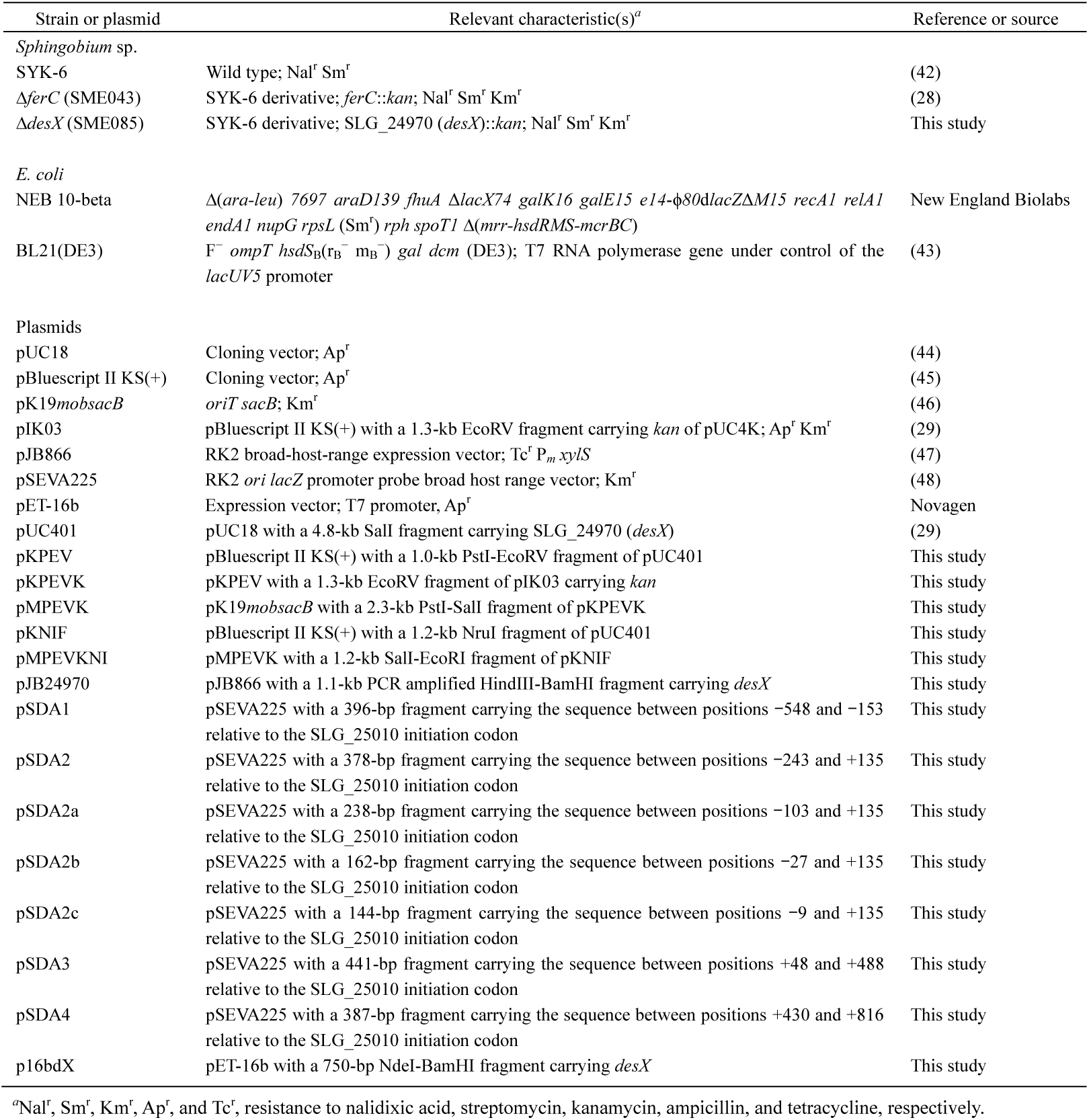
Strains and plasmids used in this study

**Fig. 5.**
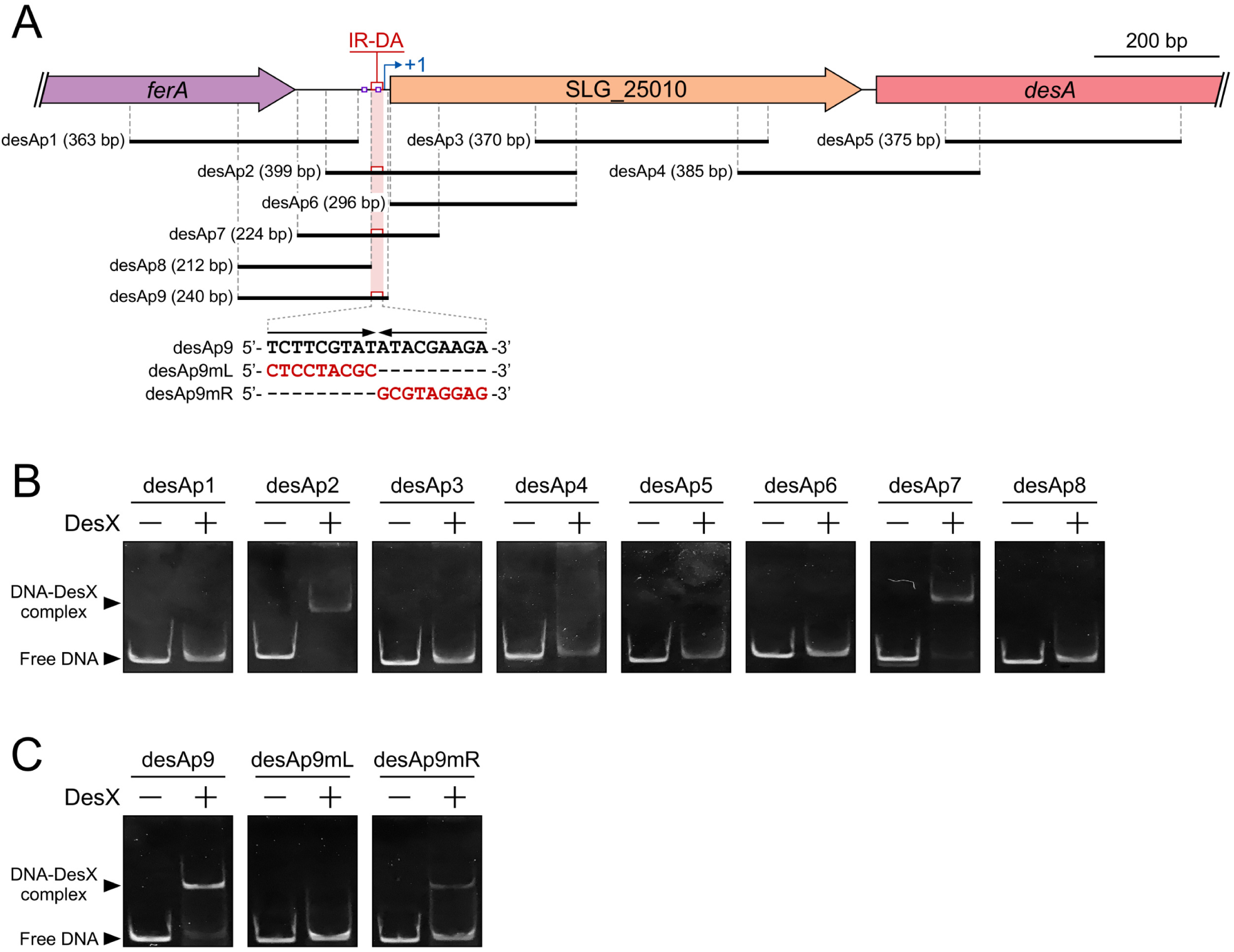
Determination of the binding region of DesX. (A) The DNA fragments used for EMSA. The transcription start site of the SLG_25010-*desA* operon is shown by a bent arrow. Putative −35 and −10 sequences and positions of IR-DA are shown by violet and red boxes, respectively. Mutated sequences of IR-DA in the desA9mL and desA9mR probes are shown. (B) EMSAs of the binding of DesX to the desAp1−desAp8 probes. The DNA probes (400 pM) were incubated in the presence (+) and absence (−) of purified DesX (8 ng protein/μl). (C) EMSAs of the binding of DesX to the desAp9 and the IR-DA mutated probes (desAp9mL and desAp9mR). Each probe (400 pM) was incubated in the presence (+) and absence (−) of purified DesX (8 ng protein/μl).

### Identification of effector molecules of DesX

Previously, we demonstrated that SA and VA are inducers of *desA* (27). Here, we also show that the transcriptions of *desA* and SLG_25010 are significantly induced in the presence of SA or VA (Fig. 3A and E). To examine the effect of SA and VA on the DNA binding of DesX, we performed EMSAs using DesX and the desAp2 probe in the presence of either SA or VA (Fig. 6). In the presence of SA, band shifts decreased in a concentration-dependent manner and disappeared entirely at 5 mM SA. In the presence of VA, a similar decrease in band shifts was observed with the desAp2 probe. Therefore, both SA and VA are effector molecules for DesX, and SA and VA have different affinities for DesX, since SA affected DNA binding of DesX at lower concentrations (5 mM SA vs. 50 mM VA) (Fig. 6).

**Fig. 6.**
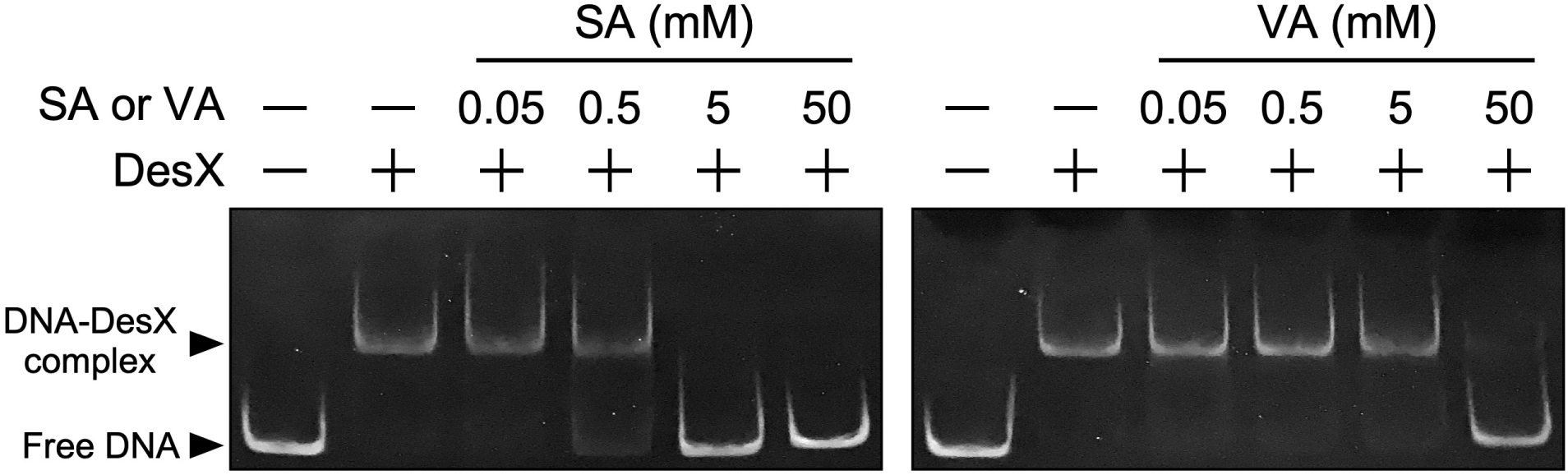
Identification of the effector molecules of DesX. EMSAs of the binding of DesX to the desAp2 probe containing IR-DA in the presence of VA or SA. Purified DesX (4 ng protein/μl) and the desAp2 probe (400 pM) were incubated in the presence (0.05, 0.5, 5, or 50 mM) or absence of SA or VA.

## DISCUSSION

This study reveals that the *ferB-ferA*-SLG_25010-*desA* gene cluster consists of the *ferBA* operon and the SLG_25010-*desA* operon, and the transcription start site of the SLG_25010-*desA* operon is located 10 bp upstream of the SLG_25010 initiation codon (Fig. S3 and S5). DesX is the only regulator that controls the transcription of the SLG_25010-*desA* operon (Fig. 3A and E); it is not involved in the transcriptional regulation of *ligM* and *desB* (Fig. 3B, C, and S8). DesX recognizes both SA and VA as effector molecules and has a higher affinity for SA than VA (Fig. 6). Based on these findings and those in our previous study (26, 27), the catabolism of SA in SYK-6 is as follows (Fig. 1): The binding of SA to DesX first triggers the derepression of *desA*, enabling the conversion of SA to 3MGA by DesA. The binding of SA to DesR also derepresses *ligM* and *desB* (27), allowing LigM to O demethylate 3MGA to GA. The aromatic ring of GA is then cleaved by DesB. Because GA is an effector molecule of LigR (26), GA binding to LigR activates the transcription of the PCA 4,5-cleavage pathway genes involved in the catabolism of the ring-cleavage product of GA.

Both DesX and DesR recognize VA and SA as effector molecules. This is reasonable for DesR because it is involved in the regulation of *ligM*, which is involved in the O demethylation of both VA and 3MGA (an intermediate metabolite of SA). Mutant analysis indicated that DesA, which is mainly involved in the O demethylation of SA, is also partially involved in the conversion of VA (29, 30). Accordingly, DesX recognizes VA as an effector molecule in addition to SA, which may enable a more efficient conversion of VA. Although DesX has a higher affinity for SA than for VA *in vitro* (Fig. 6), *desA* is transcribed at higher rate *in vivo* under VA-inducing conditions compared to SA-inducing conditions (Fig. 3A). The reason for this discrepancy is still unclear, however, the *in vivo* results may not completely reflect *in vitro* results, as the former is affected by other factors, such as the level of substrate uptake.

The transcription start site of the SLG_25010-*desA* operon is located 10 bp upstream from the initiation codon of SLG_25010, and a weak Shine-Dalgarno sequence is 8- to 6-bp upstream of the initiation codon (Fig. S5). According to the RBS calculator (31), the translation initiation rate of SLG_25010 from the putative mRNA sequence of the SLG_25010-*desA* transcript is 1.77, a rate that is markedly lower than those for *desA* (2395), *ferB* (2577), *ferA* (230), *ligM* (177), and *desB* (6023). Therefore, SLG_25010 may be poorly translated from this mRNA. Read-through from the *ferB* promoter was observed in the intergenic region between *ferA* and SLG_25010 under SA-, VA-, or FA-inducing conditions (Fig. S3C to E). Therefore, the mRNA generated from this read-through may be required for the translation of SLG_25010 (translation initiation rate, 82). The function of SLG_25010, currently annotated as a hydrolase, has not been clarified. In SYK-6 cells, a metabolite in the SA catabolism, 4-carboxy-2-hydroxy-6-methoxy-6-oxohexa-2,4-dienoate (CHMOD), is spontaneously converted to 2-pyrone-4,6-dicarboxylate (PDC). Simultaneously, CHMOD is converted to OMA by a hydrolase whose gene has not yet been identified (Fig. 1) (32). Recently, the methylesterase (DesC) and the *cis-trans* isomerase (DesD) genes were reported to be involved in the conversion of CHMOD to OMA during SA catabolism in *Novosphingobium aromaticivorans* DSM12444 (23). In the SYK-6 genome, the SLG_12720 and SLG_07230 amino acid sequences are 45% and 40% similar to those of *desC* and *desD*, respectively. However, there is no SLG_25010 ortholog in the DSM 12444 genome. It will be necessary to investigate the involvement of these genes in the conversion of CHMOD in SYK-6 in the future.

In *N. aromaticivorans* DSM 12444, SA is converted to 3MGA by the Saro_2404 gene product (DesA_NA_), whose amino acid sequence is 71% similar to that of SYK-6 DesA. The resulting 3MGA is metabolized via CHMOD as described above (23). In the DSM 12444 genome, a BLAST search reveals Saro_2407, encoding a product with 51% amino acid sequence identity with DesX. Because Saro_2404 (*desA*_NA_) and Saro_2407 (*desX* ortholog) are closely located (Fig. S9), *desA*_NA_ is probably regulated by the Saro_2407 gene product. In DSM 12444, VA is converted to PCA by the Saro_2861 gene product (LigM_NA_), which is 78% similar to the SYK-6 LigM, and PCA is metabolized via the PCA 4,5-cleavage pathway (23). DSM 12444 also possesses Saro_0803, which is 51% similar to SYK-6 DesR (Fig. S9), implying that Saro_2861 (*ligM*_NA_) is regulated by the Saro_0803 gene product (DesR ortholog) in DSM 12444.

ITTRs are known to include both activators and repressors (33). Many ITTRs that have been reported as activators are thought to bind to the upstream region of the −35 sequence on the target promoter, thus recruiting RNA polymerase (RNAP) (34, 35). In contrast, the binding sequences of the repressors HmgR and IphR are located downstream of the −10 sequence on the target promoter (36, 37). EMSAs show DesX binding to IR-DA, an inverted repeat sequence that overlaps the −10 sequence of the SLG_25010 promoter (Fig. 5). Therefore, DesX appears to repress the transcription of the SLG_25010-*desA* operon by preventing the binding of RNAP to the SLG_25010 promoter.

In conclusion, we have clarified the transcriptional regulatory system for the SA *O*-demethylase gene *desA* in SYK-6, a model bacterial degrader of lignin-derived aromatic compounds. Our present results, combined with our previous findings, have enabled us to provide an overall picture of the regulatory systems for SA and VA catabolism. This information is essential for creating engineered bacteria that can efficiently produce value-added chemicals from lignin.

## MATERIALS AND METHODS

### Bacterial strains, plasmids, culture conditions, primers, and chemicals

The bacterial strains and plasmids used in this study are listed in Table 1, and PCR primers are listed in Table 2. *Sphingobium* sp. SYK-6 and its mutants were grown at 30°C with shaking (160 rpm) in Lysogeny broth (LB), Wx medium containing SEMP, 5 mM SA, VA, or FA, and Wx-SEMP containing 5 mM SA or VA. Whenever necessary, the media for SYK-6 and its mutants and transformants were supplemented with 50 mg of kanamycin/liter, 12.5 mg of tetracycline/liter, or 12.5 mg of nalidixic acid/liter. *Escherichia coli* NEB 10-beta was used for the cloning experiments. *E. coli* BL21(DE3) was used for the expression of *desX. E. coli* strains were grown at 30°C or 37°C with shaking (160 rpm) in LB. The media for *E. coli* transformants were supplemented with 100 mg of ampicillin/liter, 25 mg of kanamycin/liter, or 12.5 mg of tetracycline/liter. SA, VA, and FA were purchased from Tokyo Chemical Industry Co. Ltd.

**Table 2.**
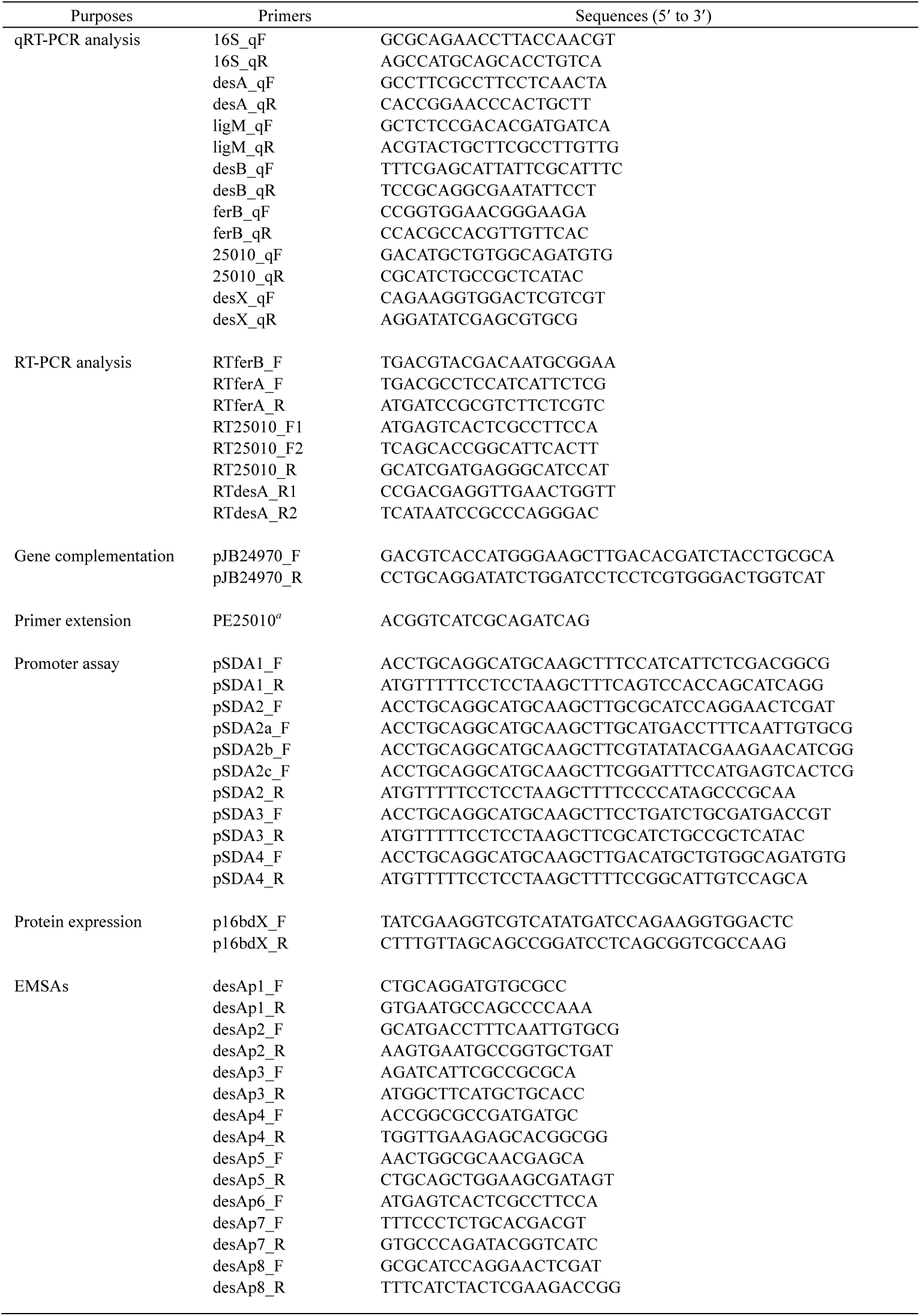
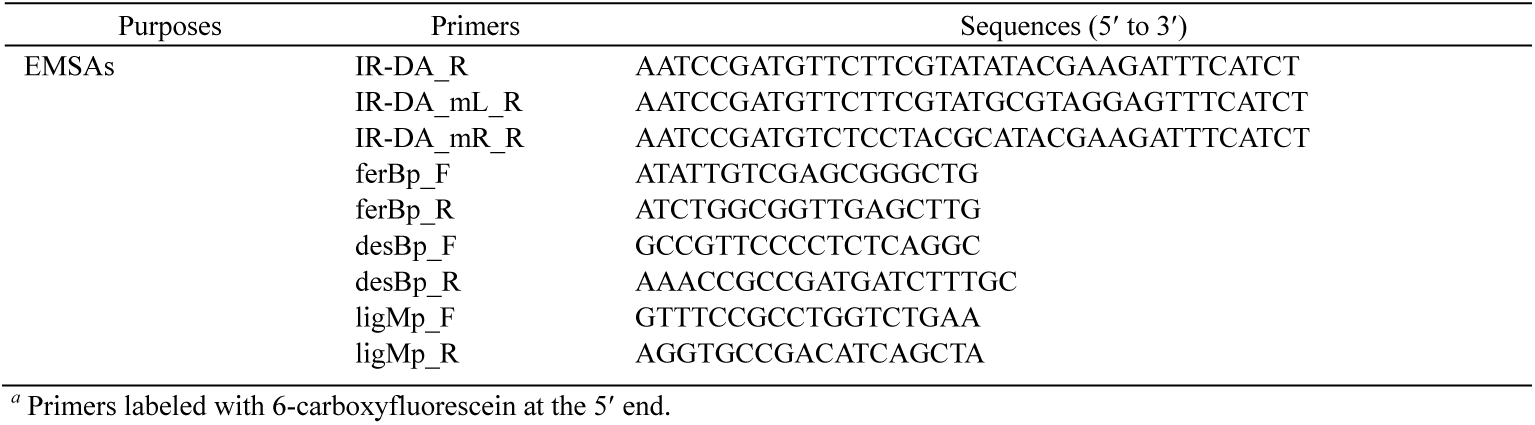
Primers used in this study

### EMSA

DNA fragments were amplified from the SYK-6 genomic DNA by PCR using the specific primers listed in Table 2. EMSAs were performed on cell extracts using a DIG Gel Shift Kit (2nd generation) (Roche). DNA fragments were labeled at their 3’ end with digoxigenin-11-ddUTP using terminal transferase. Cell extracts of SYK-6 and mutants were prepared as described previously (27). The DNA-protein binding reactions were conducted at 20°C for 20 min in a final volume of 10 μl. EMSA reaction mixtures contained cell extracts (0.4 μg protein/μl), 5 fmol digoxigenin-labeled probe, 1 μg poly-[d(I-C)], and binding buffer [20 mM HEPES, 1 mM EDTA, 10 mM (NH_4_)_2_SO_4_, 1 mM dithiothreitol, 0.2% (w/v) Tween 20, 30 mM KCl, pH 7.6]. EMSA reaction mixtures using purified DesX contained purified DesX [0.5−16 ng protein/μl], 4 fmol non-labeled probe, and binding buffer. To examine the association of DesX with effector molecules, 1 μl of a solution of SA or VA (final concentrations, 0.05, 0.5, 5, or 50 mM) was added to the reaction mixture. After incubation, 2.5 μl loading buffer [60% (vol/vol) 0.25× TBE buffer (89 mM Tris, 89 mM boric acid, and 20 mM EDTA, pH 8.0), 40% (vol/vol) glycerol, and 0.2% (vol/vol) bromophenol blue] was added, and samples were separated on a 5% native polyacrylamide gel. After electrophoresis, the labeled DNAs were detected using a CSPD-based chemiluminescence detection system (Roche) (26), and the non-labeled DNAs in the gel were stained with SYBR Gold Nucleic Gel Stain (Invitrogen) and photographed under a 470-nm blue LED.

### Construction of the SLG_24970 mutant

The 1.0-kb PstI-EcoRV fragment carrying the upstream region of SLG_24970 from pUC401 was inserted into the corresponding sites of pBluescript II KS+ to obtain pKPEV. The 1.3-kb EcoRV fragment carrying the kanamycin resistance gene (*kan*) of pIK03 was inserted into the EcoRV site of pKPEV to form pKPEVK. The 2.3-kb PstI-SalI fragment of pKPEVK was then inserted into the corresponding sites of pK19*mobsacB* to obtain pMPEVK. The 1.2-kb NruI fragment carrying the downstream region of SLG_24970 from pUC401 was cloned into the EcoRV site of pBluescript II KS+, and the 1.2-kb SalI-EcoRI fragment (including the 1.2-kb NruI fragment) from the resulting plasmid (pKNIF) was inserted into the corresponding site of pMPEVK, generating an SLG_24970 disruption plasmid, pMPEVKNI. pMPEVKNI was introduced into SYK-6 cells by electroporation, and the mutant was selected as previously described (38, 39). Disruption of the gene was confirmed by Southern hybridization analysis using the total DNA of mutant candidates digested with SalI and the digoxigenin-labeled probes (Roche), the 1.4-kb ApaI fragment carrying SLG_24970 and the 1.3-kb EcoRV fragment carrying *kan*.

The complementary plasmid, pJB24970, was constructed by amplifying a 1.1-kb fragment carrying SLG_24970 from SYK-6 genomic DNA using the primer pair listed in Table 2. The resulting fragment was inserted into the HindIII-BamHI sites of pJB866 by an NEBuilder HiFi DNA Assembly Cloning Kit (New England Biolabs). The nucleotide sequence of the insert was confirmed by sequencing. pJB866 and pJB24970 were independently introduced into Δ*24970* and SYK-6 cells by electroporation, and the growth of transformants was measured as described in the following section.

### Bacterial growth measurement

Cells of SYK-6, its mutants, and complemented strains were grown in LB for 24 h, harvested by centrifugation (5,000 × *g*, 5 min, 4°C), washed twice with Wx medium, and resuspended in 1 ml of the same medium. Cells were then inoculated into 5 ml of Wx medium containing 5 mM SA or VA to an OD_660_ of 0.2. Cells were incubated at 30°C with shaking (60 rpm) and cell growth was periodically monitored by measuring the OD_660_ using a TVS062CA bio-photorecorder (Advantec Co., Ltd.). Complemented strains of Δ*24970* were analyzed by growing cells in Wx medium containing tetracycline and 1 mM *m*-toluate, an inducer of the P_*m*_ promoter in pJB866.

### Isolation of total RNA

Cells of SYK-6 and Δ*desX* were grown in LB for 24 h, harvested by centrifugation (5,000 × *g*, 5 min, 4°C), washed twice with Wx medium, and resuspended in the same medium. The resulting cells were inoculated into 10 ml of Wx-SEMP, Wx-5 mM SA, Wx-5 mM VA, and Wx-5 mM FA to an OD_600_ of 0.2 and incubated until the OD_600_ of the culture reached 0.5−0.6. Total RNAs were isolated as described previously (27).

### qRT-PCR and RT-PCR

cDNAs were synthesized by reverse transcription using a PrimeScript II 1st Strand cDNA Synthesis Kit (Takara Bio). Total RNA (1 μg) was reverse transcribed using PrimeScript reverse transcriptase with random primers (6-mers). Reactions for each sample included a reverse transcriptase negative control to verify no genomic DNA contamination had occurred. The synthesized cDNA was purified using a NucleoSpin Gel and PCR Clean-up Kit (Takara Bio) and eluted with 30 μl of 5 mM Tris-HCl buffer (pH 8.5). qRT-PCR reactions used a THUNDERBIRD SYBR qPCR Mix (TOYOBO) and were amplified on a LightCycler 480 System II (Roche). The specific primer pairs used for qRT-PCR analyses were designed using the Primer Express version 2.0 software program (Applied Biosystems) (Table 2). qRT-PCR reaction contained 2 μl of cDNA sample, gene-specific primers (10 pmol), and THUNDERBIRD SYBR qPCR Mix (10 μl) in a total reaction volume of 20 μl. Thermal cycling conditions were as follows: 20 s at 95°C followed by 40 repeats of 3 s at 95°C and 30 s at 60°C. The LightCycler 480 System II detected and analyzed changes in the fluorescence emission and performed a melting curve analysis at the end of qRT-PCR to verify the specificity of the amplification. The amount of RNA in each sample was normalized relative to 16S rRNA, which was used as an internal standard. Each measurement was conducted in triplicate, and the means and standard deviations were calculated. RT-PCR was performed using 1 μl of cDNA sample, specific primers (Table 2), and Q5 Hot Start High-Fidelity DNA Polymerase (New England Biolabs). PCR products were electrophoresed on a 0.8% agarose gel.

### Promoter assay

Reporter plasmids were constructed using DNA fragments containing *desA* and its upstream region, which were PCR-amplified from SYK-6 total DNA and the primer pair listed in Table 2. The PCR products were inserted into the HindIII site upstream of the promoter-less *lacZ* of pSEVA225. The nucleotide sequence of the insert was confirmed by sequencing. Each plasmid was introduced into SYK-6 cells by electroporation, and SYK-6 transformants were grown for 24 h in LB containing kanamycin, harvested by centrifugation, washed twice with Wx medium, and resuspended in the same medium. The resulting cells were inoculated into 10 ml Wx-SEMP, Wx-5 mM SA, and Wx-5 mM VA containing kanamycin to an OD_600_ of 0.2 and incubated until the OD_600_ of the culture reached 0.5−0.6. Cells were harvested by centrifugation, washed twice with Wx medium, and resuspended in the same medium to an OD_600_ of 2.0. The β-galactosidase activity of the cells was measured using 2-nitrophenyl-β-D-galactopyranoside according to a modified Miller assay [https://openwetware.org/wiki/Beta-Galactosidase_Assay_(A_better_Miller)] and is expressed as Miller units as previously described (40).

### Primer extension

Total RNA was isolated from SYK-6 cells grown in Wx-5 mM SA. cDNA was synthesized from total RNA (5 μg) using a PrimeScript II 1st Strand cDNA Synthesis Kit and a 6-carboxyfluorescein-labeled PE25010 primer (2 pmol). The extended products were purified by a NucleoSpin Gel and PCR Clean-up Kit and then analyzed using an ABI 3730xl DNA analyzer (Applied Biosystems) at Macrogen.

### Expression of *desX* in *E. coli* and purification of DesX

A DNA fragment carrying *desX* was amplified by PCR using SYK-6 total DNA and the primer pair listed in Table 2. The PCR product was cloned into the NdeI site of pET-16b by an NEBuilder HiFi DNA Assembly Cloning Kit and the nucleotide sequence of the insert of the resulting p16bdX was confirmed by sequencing. *E. coli* BL21(DE3) cells harboring p16bdX were grown for 12 h at 37°C in LB containing ampicillin, and the culture was inoculated into the same fresh medium (final concentration, 1%). When the OD_600_ of the culture reached 0.5−0.6, the expression of *desX* with an N-terminal His-tag was induced for 4 h at 30°C by adding 1 mM isopropyl-β-D-thiogalactopyranoside. The cells were harvested by centrifugation (5,000 × *g*, 5 min, 4°C), washed twice with 50 mM Tris-HCl buffer (pH 7.5) containing 300 mM NaCl and 100 mM imidazole (buffer A), resuspended in buffer A, and then disrupted using an ultrasonic disintegrator. The supernatant obtained by centrifugation (19,000 × *g*, 15 min, 4°C) was applied to a His SpinTrap column (GE Healthcare) previously equilibrated with buffer A. After centrifugation (100 × *g*, 1 min, 4°C), samples were washed three times with buffer A, and then His-tag-fused DesX was eluted with 50 mM Tris-HCl buffer (pH 7.5) containing 300 mM NaCl and 500 mM imidazole. Purified DesX was desalted and concentrated by centrifugal filtration using an Amicon Ultra centrifugal filter unit (10 kDa cutoff; Merck Millipore). Before the assay, insoluble aggregates in the DesX solution were removed by centrifugal filtration using an Ultrafree-MC filter (Merck Millipore). The protein concentration was determined by the Bradford method (41) with bovine serum albumin as the standard (Nacalai tesque). The gene expression and purity of the preparation were examined by SDS-12% PAGE, and the protein bands in gels were stained with Coomassie Brilliant Blue.

## ACKNOWLEDGMENTS

This work was supported in part by Grant-in-Aid for JSPS Fellows 16J11003 and Research grant 201916 of the Forestry and Forest Products Research Institute.

